# Atorvastatin effectively inhibits late replicative cycle steps of SARS-CoV-2 *in vitro*

**DOI:** 10.1101/2021.03.01.433498

**Authors:** María I. Zapata-Cardona, Lizdany Flórez-Álvarez, Wildeman Zapata-Builes, Ariadna L. Guerra-Sandoval, Carlos M. Guerra-Almonacid, Jaime Hincapié-García, María T. Rugeles, Juan C. Hernandez

## Abstract

**Introduction:** SARS-CoV-2 (Severe Acute Respiratory Syndrome Coronavirus 2) has caused a pandemic of historic proportions and continues to spread worldwide. Currently, there is no effective therapy against this virus. This article evaluated the *in vitro* antiviral effect of Atorvastatin against SARS-CoV-2 and also identified the interaction affinity between Atorvastatin and three SARS-CoV-2 proteins, using *in silico* structure-based molecular docking approach.

**Materials and methods:** The antiviral activity of Atorvastatin against SARS-CoV-2 was evaluated by three different treatment strategies using a clinical isolate of SARS-CoV-2. The interaction of Atorvastatin with Spike, RNA-dependent RNA polymerase (RdRp) and 3C-like protease (3CLpro) was evaluated by molecular docking.

**Results:** Atorvastatin showed anti-SARS-CoV-2 activity of 79%, 54.8%, 22.6% and 25% at 31.2, 15.6, 7.9, and 3.9 µM, respectively, by pre-post-treatment strategy. In addition, atorvastatin demonstrated an antiviral effect of 26.9% at 31.2 µM by pre-infection treatment. This compound also inhibited SARS-CoV-2 in 66.9%, 75%, 27.9% and 29.2% at concentrations of 31.2, 15.6, 7.9, and 3.9 µM, respectively, by post-infection treatment. The interaction of atorvastatin with SARS-CoV-2 Spike, RdRp and 3CL protease yielded a binding affinity of −8.5 Kcal/mol, −6.2 Kcal/mol, and −7.5 Kcal/mol, respectively.

**Conclusion:** Our study demonstrated the *in vitro* anti-SARS-CoV-2 activity of Atorvastatin, mainly against the late steps of the viral replicative cycle. A favorable binding affinity with viral proteins by bioinformatics methods was also shown. Due to its low cost, availability, well-established safety and tolerability, and the extensive clinical experience of atorvastatin, it could prove valuable in reducing morbidity and mortality from COVID-19.

**Importance:** The COVID-19 pandemic constitutes the largest global public health crisis in a century, with enormous health and socioeconomic challenges. Therefore, it is necessary to search for specific antivirals against its causative agent (SARS-CoV-2). In this sense, the use of existing drugs may represent a useful treatment option in terms of safety, cost-effectiveness, and timeliness. Atorvastatin is widely used to prevent cardiovascular events. This compound modulates the synthesis of cholesterol, a molecule necessary in different stages of the viral replicative cycle. Our study demonstrated the antiviral potential of atorvastatin against SARS-CoV-2, using an *in vitro* model. Furthermore, the ability of Atorvastatin to directly interfere with three viral targets (Spike, RdRp and 3CL protease) was demonstrated by bioinformatic methods. This compound is a well-studied, low-cost, and generally well-tolerated drug, so it could be a promising antiviral for the treatment of COVID-19.

## Introduction

COVID-19 (coronavirus disease 2019) is a disease caused by the SARS-CoV-2 virus (Severe Acute Respiratory Syndrome Coronavirus 2) reported for the first time in Wuhan, China [1]. On March 12, 2020, the WHO (World Health Organization) declared COVID-19 a pandemic [2]. Since then, it has affected 218 countries and territories worldwide, causing enormous human health consequences [1,3].

Currently, disease control has been based on symptom management, including the use of convalescent plasma, synthetic antibodies, interferon, low-dose corticosteroids, IL-1 and IL-16 inhibitors, Azithromycin, Remdesivir, Baricitinib, Lopinavir/Ritonavir, Hydroxychloroquine, and, in severe cases, supportive care (oxygen and mechanical ventilation) [4, 5]. Although there may be approved drugs [6], there is still no conclusive information on their effectiveness and they have limited uses. For this reason, the search for effective and specific anti-SARS-CoV-2 antivirals is necessary [7]. The development of new drugs involves the evaluation of the pharmacokinetics and pharmacodynamic safety, with the implementation of large-scale production and distribution, a process that takes many years. Due to this panorama and the rapid expansion of the pandemic, the use of existing drugs may represent a useful COVID-19 treatment option in terms of safety, cost-effectiveness, and timeliness [8].

Atorvastatin (ATV) belongs to statins, a group of hypolipidemic drugs that acts inhibiting HMG-CoA reductase. ATV was approved by the FDA (Food and Drug Administration) and EMA (European Medicines Agency) to prevent cardiovascular events in patients with cardiac risk factors or with abnormal lipid profiles [9, 10]. The ability of ATV to modulate cholesterol synthesis has previously been related to antiviral activity against viruses such as Hepatitis C virus [11], Dengue virus [12], Zika virus [13], Influenza A virus [14] and human parainfluenza virus type 1 [15]. This drug also participates in modulating several cellular metabolic pathways that could affect the viral replicative cycle [11, 12].

It has been proposed that statins could be an effective therapeutic strategy against SARS-CoV-2 infection [16]. Recently, it has been reported that statin treatment was associated with a reduced risk of mortality in patients diagnosed with COVID-19 [17, 18]. Furthermore, during *in silico* analysis, ATV interacts with the 3CL protease of SARS-CoV-2 [19]. However, there is no *in vitro* evidence of its antiviral effect against SARS-CoV-2 or its binding affinity for other viral proteins such as Spike and RNA-dependent RNA polymerase (RdRp). This article evaluated the *in vitro* antiviral effect of the ATV against SARS-CoV-2, and identified the interaction affinity between ATV and three SARS-CoV-2 proteins, using an *in silico* structure-based molecular docking approach.

## Results

### Atorvastatin showed a low cytotoxicity on Vero E6

Before the antiviral activity evaluation, the cytotoxic effect of ATV on Vero E6 cells was determined by MTT assay. As shown in **Figure 1**, the Vero E6 viability was higher than 90% at a concentration of 31.2 µM or less of ATV. At higher ATV concentrations (62.5 to 250 µM), cell viability was lower or equal to 48.4%. The CC50 calculated for ATV was 50.3 µM.

**Figure 1.**
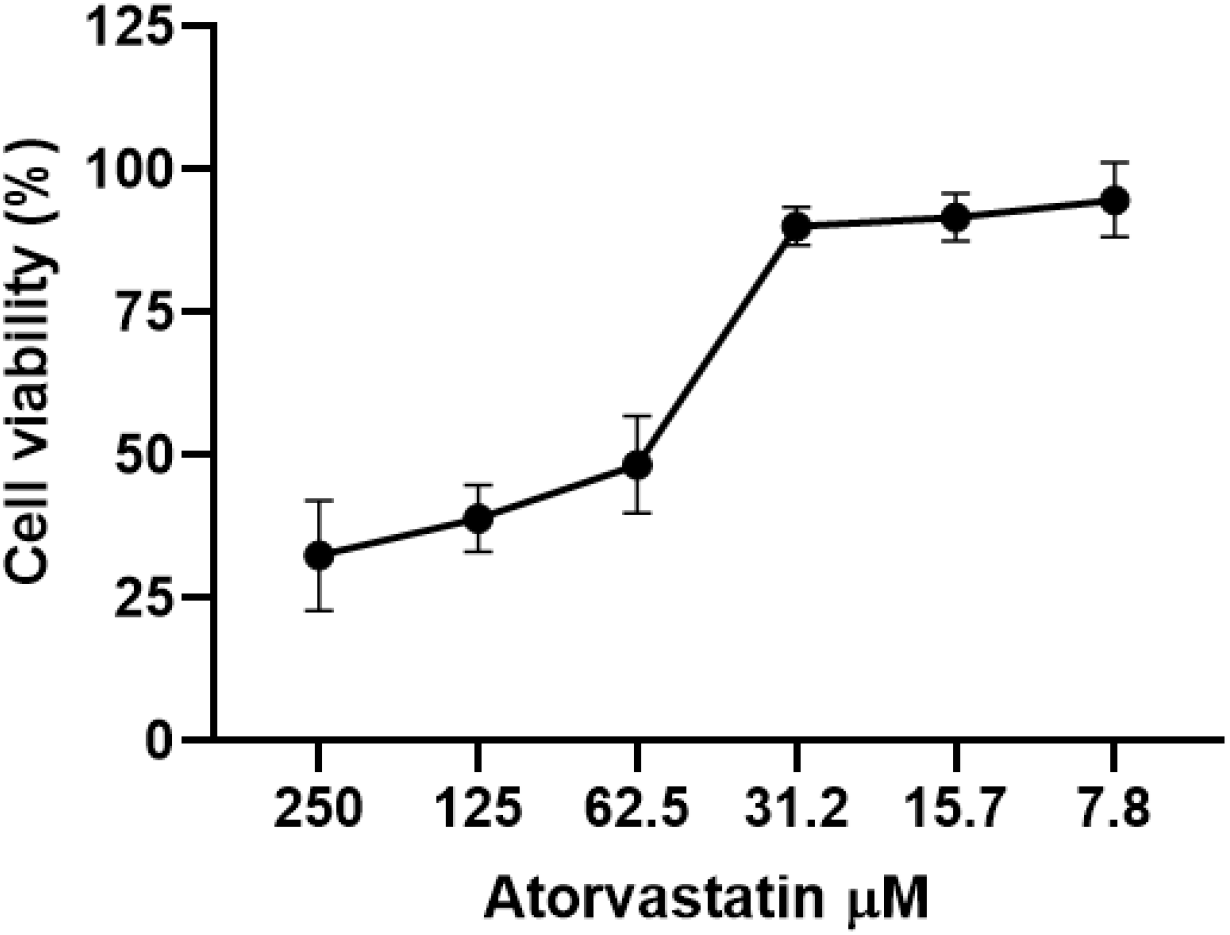
Atorvastatin showed a low cytotoxicity on Vero E6. Viability of Vero E6 after 48h of atorvastatin treatment (from 7.8 to 250 µM). Data were presented as Mean ± SEM. The viability percentages of the treated cell were calculated based on untreated control. Two independent experiments with four replicates each were performed (n=8).

Cell viability was not affected by CQ and Heparin (positive controls of viral inhibition) at the concentrations that were evaluated **(Supplementary figure 1).** The CC50 obtained by CQ was more than 100 µM, and by Heparin was more than 100 µg/mL.

### ATV exhibited antiviral effects against SARS-CoV-2 in a dose-dependent manner

To evaluate the antiviral activity of ATV against SARS-CoV-2, a pre-post treatment strategy was performed in Vero E6 cells. ATV showed inhibition percentages of SARS-CoV-2 of 79% (p=0.002), 54.8% (p=0.002), 22.6% (p=0.04) and 25% (p=0.03) at concentrations of 31.2, 15.6, 7.9, and 3.9 µM, respectively **(Figure 2).** Based on these data, the EC50 calculated for ATV was 15.4 µM, with a selectivity index of 3.3.

**Figure 2.**
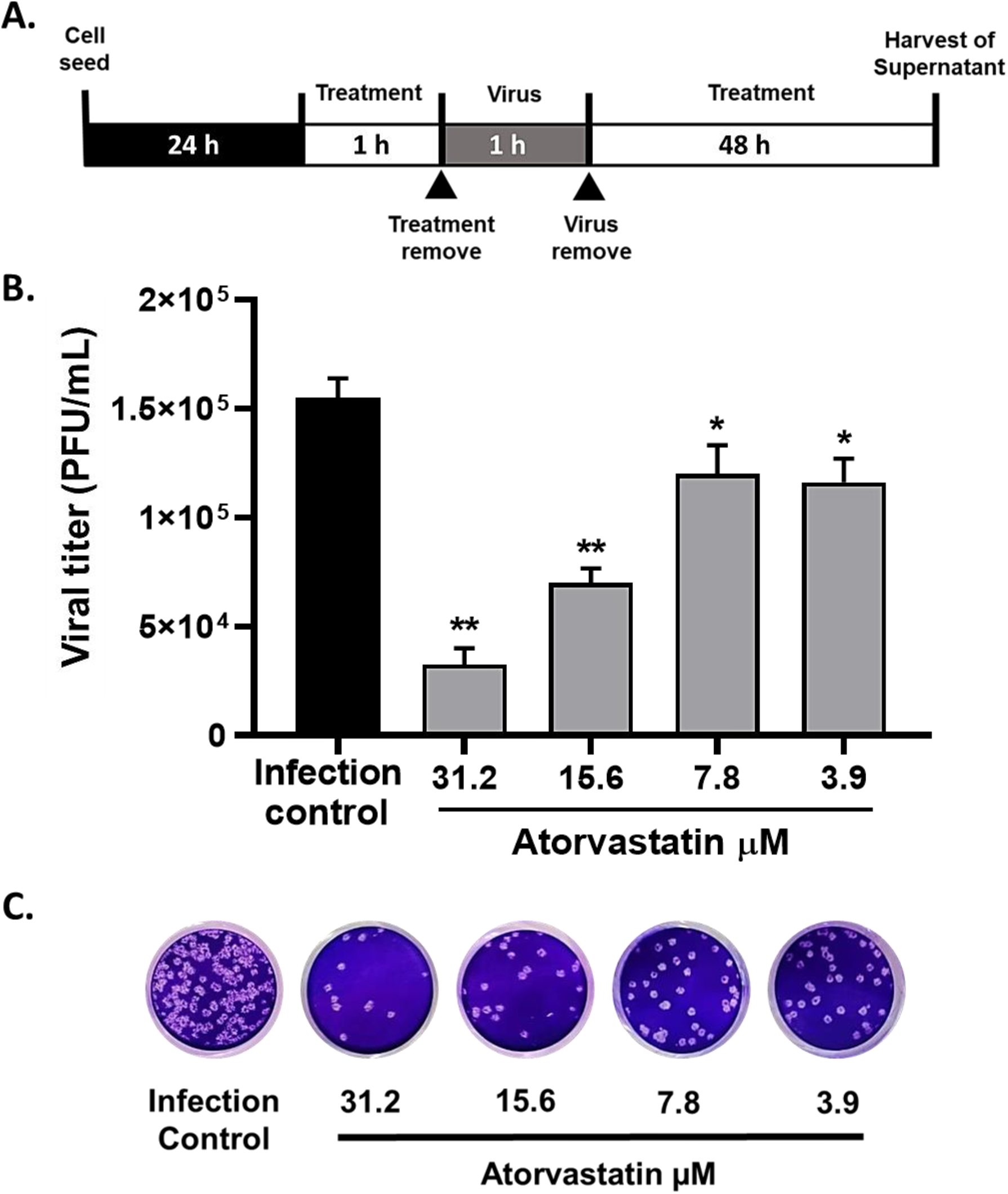
Atorvastatin exhibited an antiviral effect against SARS-CoV-2 in a dose-dependent manner. **A.** Schematic representation of the pre-post treatment strategy. **B.** Reduction of SARS-CoV-2 titer (PFU/mL) on Vero E6 supernatants after ATV treatment by pre-post treatment (n=4). Data were presented as Mean ± SEM. Mann-Whitney test *p ≤ 0.05, ** p ≤ 0.01 **C.** Representative figure of plaque assays on Vero E6 cells of pre-post-treatment of ATV against SARS-CoV-2.

Chloroquine (positive control of viral inhibition) showed inhibition percentages of 100% (p=0.002), 99.9%(p=0.002), 97.5% (p=0.002) and 55.7% (p=0.002) at 100, 50, 25, and 12.5 µM concentrations, respectively **(Supplementary figure 2).** An EC50 value of 13.5 µM was found by CQ, with a selectivity index of >7.4.

### ATV inhibited SARS-CoV-2 through post-infection treatment

Once an antiviral effect against SARS-CoV-2 was observed, the pre-infection and post-infection treatments were done independently to infer the viral replication step affected by the ATV treatment. During the pre-infection treatment, anti-SARS-CoV-2 activity was observed at 31.2 µM (inhibition of 26.9%, p=0.012). Inhibition percentages of 8.1%, 20.8% and 13.9% were seen at concentrations of 15.6, 7.9 and 3.9 µM of ATV, respectively **(Figures 3A-C).** The EC50 calculated for ATV was 12.1 µM, with a selectivity index of 4.2, by the pre-infection treatment.

**Figure 3.**
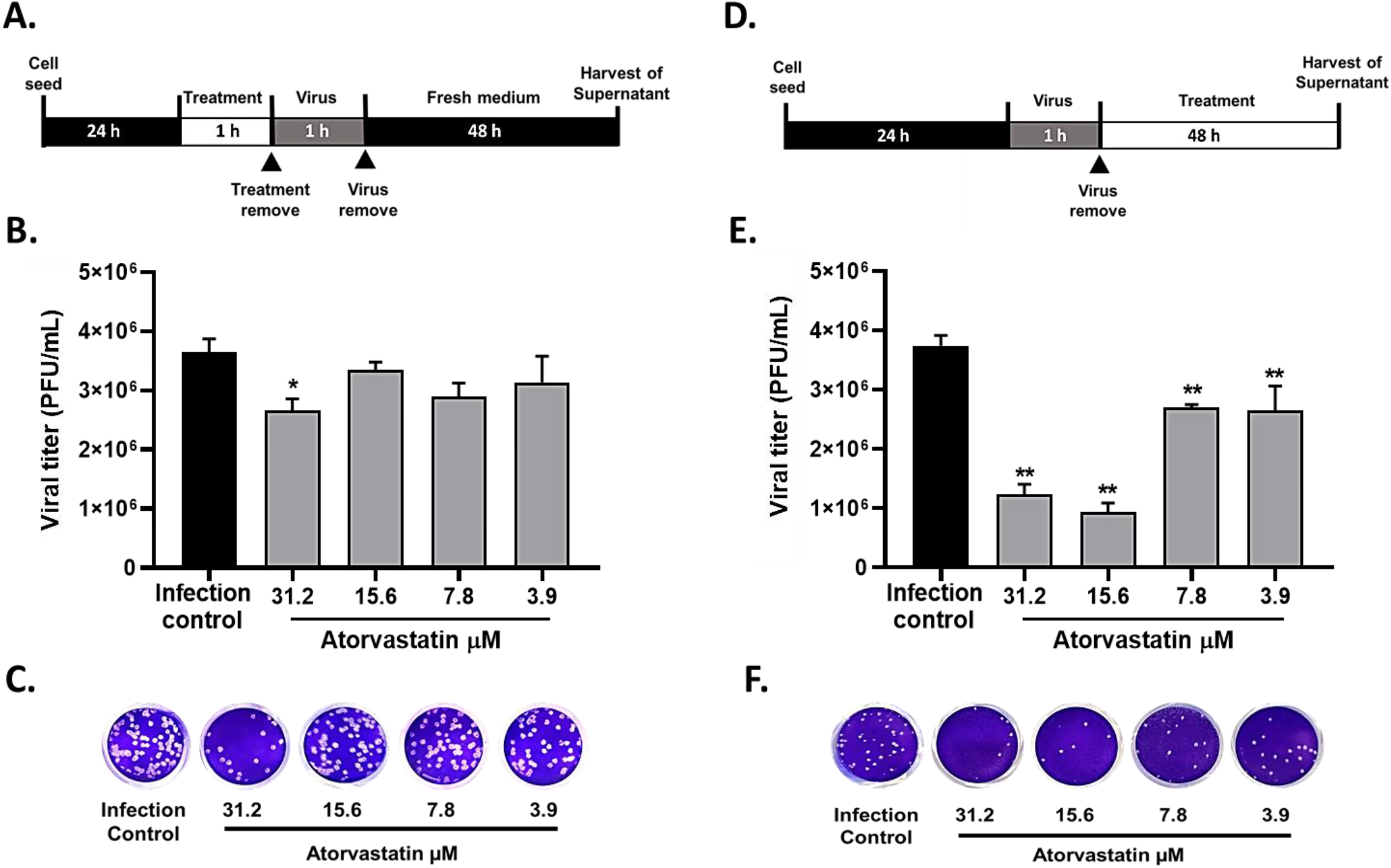
ATV inhibited SARS-CoV-2 during pre-infection and post-infection treatment. **A.** Schematic representation of the pre-infection treatment. **B.** Reduction of SARS-CoV-2 titer (PFU/mL) on Vero E6 supernatants after pre-infection treatment with ATV. Data were presented as Mean ± SEM (n=4). Mann-Whitney test *p ≤ 0.05, ** p ≤ 0.01. **C.** Representative figure of plaque assays on Vero E6 cells of pre-infection treatment of ATV against SARS-CoV-2. **D.** Schematic representation of the post-infection treatment. **E.** Reduction of SARS-CoV-2 titer (PFU/mL) on Vero E6 supernatants after post-infection treatment with ATV. **F.** Representative figure of plaque assays on Vero E6 cells of post-infection treatment with ATV against SARS-CoV-2.

In comparison, as shown in **Figures 3D-F** the viral titer of SARS-CoV-2 was significantly reduced through post-infection treatment with ATV at all concentrations evaluated. Inhibition percentages of 66.9% (p=0.004), 75.0% (p=0.004), 27.9% (p=0.004), and 29.2% (p=0.006) were obtained at concentrations of 31.2, 15.6, 7.9, and 3.9 µM of ATV, respectively (EC50= 11.1 µM, IS= 4.5).

The positive controls for viral inhibition during pre-infection and post-infection treatments were Heparin and Chloroquine, respectively. By, pre-infection treatment, Heparin inhibited SARS-CoV-2 at concentrations of 100 µg/mL (80.6%, p=0.028), 50 µg/mL (82.9%, p=0.028), 25 µg/mL (87.3%, p=0.028), and 12.5 µg/mL (79.9%, p=0.028) **(Supplementary figure 3A)**. On the other hand, as shown in **Supplementary figure 3B**, inhibition percentages of 99.2% (p=0.009), 98.3% (p=0.009), 74.8% (p=0.009) and 23.6% (p=0.033) were obtained at concentrations of 100, 50, 25, and 12.5 µM concentrations of Chloroquine, using post-infection treatment (EC50=12.76, IS>7.8).

### Binding site determination of viral proteins

Before molecular docking, pockets and expected amino acids were selected to assess the interaction affinity of SARS-CoV-2 proteins with ATV. The Spike protein (PDB:6VYB) had 79 pockets **(Supplementary figure 4A)**, the pocket score value 82.76 was the number one and consisted of 114 amino acids. The 3CL protease (PDB:6M2N) showed 15 pockets **(Supplementary figure 4B)**, and the highest score was obtained in pocket number one and consisted of 19 amino acids. Finally, the SARS-CoV-2 RdRp (PDB:6M71) had 37 pockets **(Supplementary figure 4C)**. The pocket with the best score was the number one with 8.79 and consisted of 23 amino acids. The first binding site is the highest pocket score for all pockets in each protein analyzed, favoring the interaction and binding with different ligands. The grid centers of these pocket sites are shown in **table 1**.

**Table 1:**
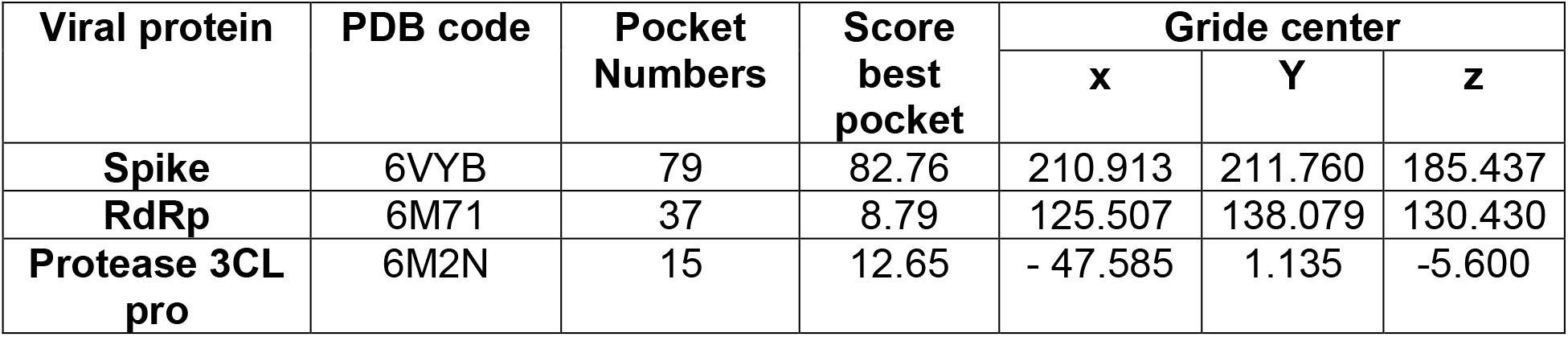
**PrankWeb result summary of SARS-CoV-2 proteins.**

| Viral protein | PDB code | Pocket Numbers | Score best pocket | Gride center |  |  |
| --- | --- | --- | --- | --- | --- | --- |
|  |  |  |  | x | Y | z |
| **Spike** | 6VYB | 79 | 82.76 | 210.913 | 211.760 | 185.437 |
| **RdRp** | 6M71 | 37 | 8.79 | 125.507 | 138.079 | 130.430 |
| **Protease 3CL pro** | 6M2N | 15 | 12.65 | - 47.585 | 1.135 | -5.600 |

### ATV demonstrated favorable binding affinities with SARS-CoV-2 proteins by molecular docking

The binding affinity of all docked and analyzed compounds with their binding energy ranking is shown in **table 2**. The results revealed that ATV exhibited high interaction affinity and coupling with the different hydrophobic pockets selected from the Spike (−8.5 Kcal/mol), RdRp (−6.2 Kcal/mol) and 3CL protease (−7.5 Kcal/mol) proteins of SARS-CoV-2 (**table 2**).

**Table 2:** **Molecular docking of Atorvastatin and Chloroquine (positive control of viral inhibition) with different crystal structures of SARS-CoV-2 proteins.**

|  | Score with the pocket of viral proteins (kcal/mol) |  |  |
| --- | --- | --- | --- |
| Ligand name | Spike (PDB:6VYB) | RdRp (PDB:6M71) | Protease 3CL pro (PDB:6M2N) |
| **Atorvastatin** | -8.5 | -6.2 | -7.5 |
| **Chloroquine** | -5.9 | -5.6 | -6.3 |

The molecular interactions of ATV as a ligand with Spike, RdRp and 3Clpro proteins of SARS-CoV-2 were numerous and explained the high affinity presented **(Figure 4).** Specifically, ATV formed five distance hydrogen bonds with Spike protein (distances of 1,998 Å, 2,867 Å, 3,280 Å, 3,590 Å, 3,336 Å). In these bonds participated amino acids ASN1023, THR1027, SER1021, and GLU1017 of Spike. In addition, ATV formed hydrophobic interactions such as Pi-Alkyl and Pi-Sigma with Spike protein. Amino acid LEU1024 was involved in Pi-pi T shaped, while ALA1020 and ALA1026 participated in Pi-alkyl bonds **(Figure 4A-B).**

**Figure 4.**
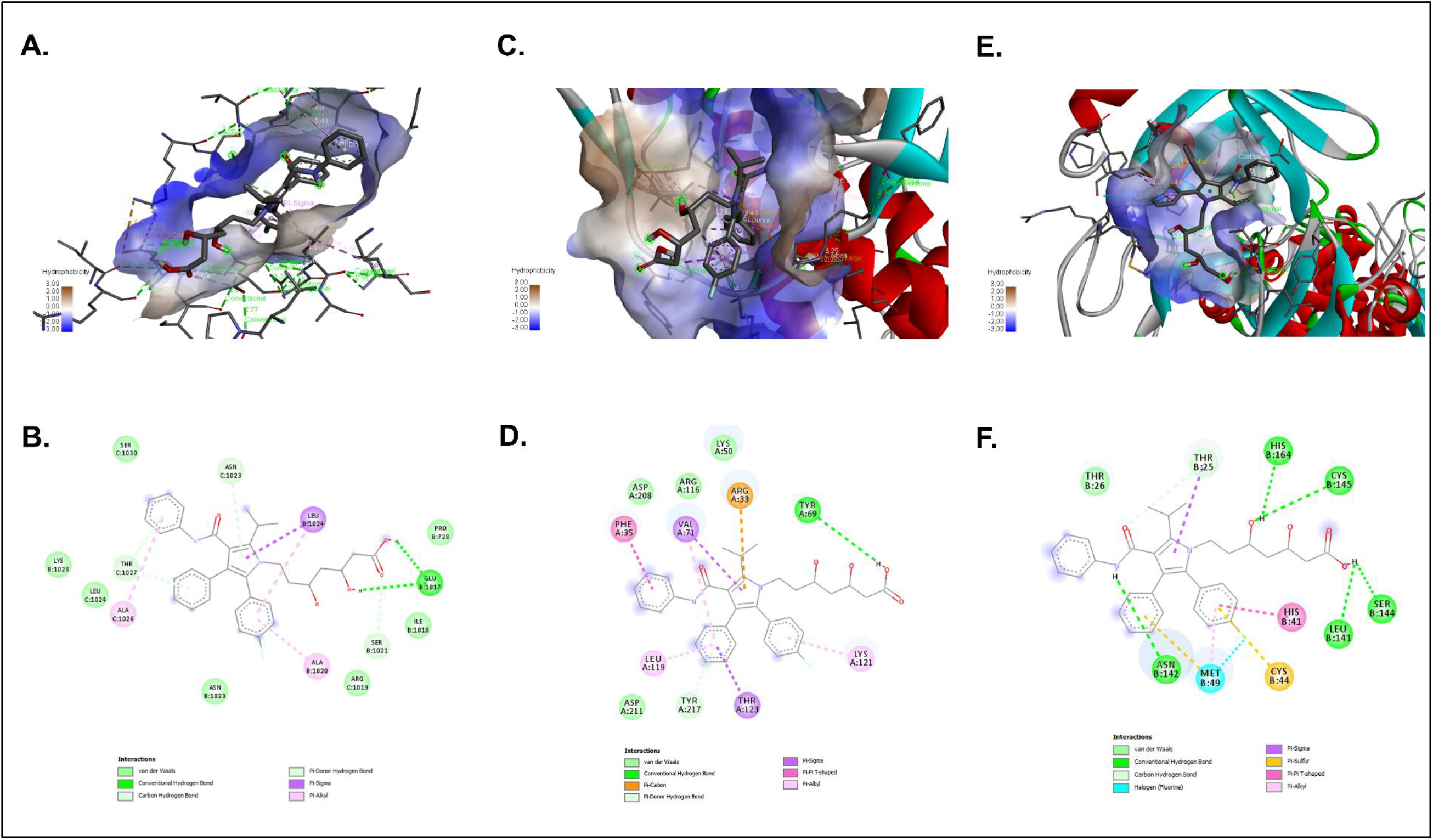
Interaction of ATV with SARS-CoV-2 proteins by molecular docking. 3D and 2D representations of the main interaction of ATV and three SARS-CoV-2 proteins by molecular docking. The images were obtained using BIOVIA Discovery Studio Visualizer 16.1. ATV interaction with: Spike (PDB:6VYB) depicted in 3D **(A)** and 2D **(B)**; RdRp (PDB:6M71) represented in 3D **(C)** and 2D **(D)**; 3CL protease (PDB: 6M2N) of SARS-CoV-2 depicted in 3D **(E)** and 2D **(F).** The interactions formed in the complexes are described in each figure.

ATV formed two conventional hydrogen bonds with the amino acids TYR69 and TYR217 of SARS-CoV-2 RdRp (Distances of 2,879 Å and 3,425 Å, respectively). Other types of interactions were also seen, such as donor hydrogen bonds that were part of the hydrophobic pocket. In the same way, hydrophobic interactions were found with the type of bonds pi-sigma, pi-alkyl, Pi-pi T shaped and Pi-cation. Amino acids THR123 and VAL71 of RdRp formed bonds pi-sigma with ATV, LEU119 and LYS121 participated in pi-alkyl bonds, PHE36 formed bonds Pi-pi T shaped and ARG33 Pi-cation interactions **(Figure 4C and 4D)**.

On the other hand, ATV formed six hydrogen bonds with HIS164, CYS145, ASN142, LEU141, SER144, and THR25 of 3CL protease, with distances of 3,121 Å, 3,197 Å, 2,637 Å, 2,477 Å, 3,024 Å, 2,944 Å, respectively. In this complex, it also participated in hydrophobic interactions such as pi-sigma, pi-alkyl and pi-sulf **(Figure 4E-F).**

Additionally, Chloroquine (positive inhibition control) showed the binding affinity of - 5.9 Kcal/mol, −5.6 Kcal/mol, and −6.3 Kcal/mol with Spike, RdRp and 3CL protease of SARS-CoV-2, respectively (**table 2**). Chloroquine formed two hydrogen bonds with Spike protein (distances 3.897 and 3.302 Å) and hydrophobic interactions such as Pi-sigma and Pi-alkyl with the amino acids LEU1024, THR1027, and ALA1026 of Spike **(Supplementary figures 5A and 5B).** Further, a hydrogen bond was formed between CQ and amino acids TYR217 of RdRp (distance of 2,544 Å). Six hydrophobic interactions (Pi-pi Stacked, Pi-sigma, Pi-Alkyl) were observed with amino acids THR123 (3,468 Å), PHE35 (5,386 Å) (5,053 Å), ARG33 (4,673 Å) and VAL71 (5,006 Å) of SARS-CoV-2 RdRp **(Supplementary figures 5C and 5D)**. Finally, Chloroquine formed two hydrogen bonds (distances of 2,959 Å and 2,822 Å) and hydrophobic interactions (Pi-stacked, Pi-sigma, Pi-alkyl) with 3CL protease **(Supplementary figures 5E and 5F)**.

## Discussion

In the present study, the anti-SARS-CoV-2 effect of Atorvastatin was identified by the treatment of Vero E6 cells, at different times of infection. ATV has been shown to inhibit HMG-CoA reductase, affecting cholesterol synthesis and the production of isoprenoid metabolites [13]. Previously, it has been reported that cholesterol-modifying drugs could exert antiviral effects by modulating cellular metabolic pathways required for the replicative cycle, or by direct effect against viral particles [20, 21]. Concerning SARS-CoV-2, it has been suggested that statins may help to reduce viral entry and transmission [22]. Based on our results, alteration in the cholesterol metabolism by ATV treatment seems to affect different stages of the SARS-CoV-2 replicative cycle where this lipid is involved, such as adhesion [23]; endocytosis [24, 25] and fusion [23]; replication [26] and viral particle assembly [27, 28].

A previous study suggested that other statin, Fluvastatin, reduced SARS-CoV-2 entry into the respiratory epithelium cells [29]. Similar to this report, our study demonstrated that ATV inhibited SARS-CoV-2 at 31.2 µM by pretreating cells for 1h. These findings suggest that the anti-SARS-CoV-2 activity of ATV affected at early stages of the viral replicative cycle. This effect could be related to the fact that the ACE2 cell receptors, used by SARS-CoV-2 to enter the cell, are present in lipid rafts, cholesterol-rich microdomains present at the cell membrane [24, 25]. In addition, a study showed that cholesterol transports ACE2 to sites where SARS-CoV-2 effectively enters the cell [23]. Therefore, an alteration at the level of cholesterol present at the cell membrane by ATV treatment, could affect viral attachment; however, additional experiments are needed to confirm this mechanism.

It has been also reported that statins upregulate ACE2 through epigenetic modifications [30]. Although this would be expected to promote viral infection, there is not current evidence indicating that statins enhance viral entry into host cells [31]. Conversely, the increase in the ACE2 receptor expression could be associated with a reduction of the severity of Acute Respiratory Distress Syndrome (ARDS) [32]. In this regard, ACE2 metabolizes the vasoconstrictor peptide angiotensin II to produce angiotensin 1-7, which is involved in reducing inflammation, tissue damage, and pulmonary edema [33]. Ongoing clinical studies would provide information on the effectiveness of ATV in preventing or mitigating these effects in patients with COVID-19 [34, 35].

Additionally, in this study, Vero E6 cells were treated with ATV for 48 h after infection, obtaining reduction of the SARS-CoV-2 titer at all evaluated concentrations. Regarding this strategy, it has been reported that statins could affect steps, such as replication, assembly and budding of infectious viral particles [15, 36–39]. This effect could be explained because the stages after viral entry of enveloped virus also depend on cholesterol biosynthetic pathways [20]. Specifically, it was reported that HMG-CoA reductase is necessary to produce lipid droplets (LD) [39]. These cell organelles have been recently proposed as hubs for SARS-CoV-2 genome replication and viral particle assembly [40, 41]. According to the above, ATV treatment could affect the replication and assembly stages of SARS-CoV-2 by blocking LD production. Furthermore, ATV could be altering the envelope cholesterol of the nascent viral particles, affecting their infectious capacity [21].

During the replicative cycle of Coronaviruses, Spike, RdRp and 3CL protease are well-characterized drug targets. Spike protein participates in the binding to host cell receptors and viral fusion to the cell membrane [42]. RdRp is involved in the nucleotide-binding and catalysis of viral RNA replication [43], and 3CL protease is the main viral protease, essential for processing polyproteins [19]. In addition to the results obtained *in vitro*, our study predicted the interactions between ATV and these SARS-CoV-2 proteins using molecular docking. ATV showed the highest binding affinity with amino acids present at the active site of Spike (−8.5 Kcal/mol) followed by 3CL protease (−7.5 Kcal/mol) and RdRp (−6.2 Kcal/mol). These binding energies were even equal to or higher than previously reported candidates for COVID -19 treatment [44, 45].

It should be kept in mind that the best candidate for a drug against SARS-CoV-2 is the molecule that can specifically bind to one of the targets mentioned above to form a thermodynamically stable complex. As shown in **Figure 4,** complexes between ATV and viral proteins were stabilized mainly by hydrogen bonds, in which ATV can act as an electron acceptor or a simultaneous donor and acceptor. Additionally, hydrophobic interactions (π-alkyl interaction, π-Sulfure, π-π Stached) also contributed to the stability of the complexes. Considering the above, ATV could be a plausible candidate against SARS-CoV-2. In fact, an *in silico* study showed that fluvastatin, lovastatin, pitavastatin and rosuvastatin may efficiently inhibit the SARS-CoV-2 protease [19].

According to our *in silico* and *in vitro* results, it could be suggested as antiviral mechanisms of ATV, the interaction with the Spike protein expressed on the SARS-CoV-2 envelope, affecting the binding of the virus with the cell receptor or affecting viral fusion. In addition, ATV could inhibit viral replication and proteolytic maturation by blocking with SARS-CoV-2 nonstructural proteins such as RdRp and 3CL protease.

Finally, some studies have shown that statin therapy is associated with a 30% reduction in fatal or severe disease in COVID-19 patients, a lower likelihood of admission to the intensive care unit [22, 46] and an increased chance of asymptomatic infection [22]. However, more studies are needed to evaluate the *in vivo* antiviral activity of ATV against SARS-CoV-2 to allow a better understanding of the benefits of ATV treatment in patients diagnosed with COVID-19, although the available evidence suggests that this therapy could be effective.

## Conclusion

Atorvastatin demonstrated potential antiviral activity against SARS-CoV-2, affecting different stages of the replicative cycle of this virus, especially at phases after viral entry. Due to low cost, availability, well-established safety and tolerability, and the extensive clinical experience of atorvastatin, it could prove valuable in reducing morbidity and mortality from COVID-19.

## Materials and methods

### Preparation of compounds

ATV was purchased from Biogen Idec, Inc. (Cambridge, MA). It was solubilized in dimethyl sulfoxide (DMSO) (Sigma-Aldrich) at a final concentration of 100 mM. Chloroquine (CQ) was bought in Sigma-Aldrich (St. Louis, MO, USA), and it was diluted to 15 mM in phosphate buffered saline (PBS, Lonza, Rockland, ME, USA). Heparin (positive inhibition control) was bought in B. Braun Melsungen AG and was diluted to 1 mg/mL in PBS. The stock solutions of ATV and CQ were frozen at −80°C, and Heparin was stored at 4 °C, until use.

### Cells and virus

Vero E6 cells from *Cercopithecus aethiops* kidney were grown in Dulbecco’s Modified Eagle Medium (DMEM, Sigma-Aldrich) supplemented with 2% heat-inactivated fetal bovine serum (FBS, Gibco), 2 mM L-glutamine (Gibco), 1% Penicillin-Streptomycin (Gibco). The incubation conditions were 37°C, 5% CO_2_ atmosphere, and relative humidity. The cells were infected with a viral stock produced from a Colombian isolate of SARS-CoV-2 (hCoV-19/Colombia/ANT-UdeA-200325-01/2020) [47]. The virus was propagated on Vero E6 cells. All experimental studies involving infectious SARS-CoV-2 were conducted within a biosafety level 3 laboratory.

### Cytotoxicity assay

The MTT (3-(4,5-dimethylthiazol-2-yl)-2,5-diphenyltetrazolium bromide) assay was performed to evaluate the cytotoxicity of ATV on Vero E6 cells. Briefly, cells were seeded in 96-well plates at a density of 1.0×10^4^ cells/well in DMEM supplemented with 2% FBS. Cultures were incubated for 24h at 37°C and 5% CO_2_. Then, 150uL/well of double serial dilutions of ATV ranging from 7.8 to 250 µM were prepared and added to each well. After 48h of incubation, the supernatants were removed, cells were washed twice with PBS and 30uL/well of MTT (2 mg/mL) were added. Plates were incubated for 2h at 37°C, with 5% CO_2_, protected from light. Then, 100uL/well of DMSO were added to dissolve the formazan crystals. Optical density (OD) was recorded at 570 nm using a Multiskan GO spectrophotometer (Thermo). Cell viability was calculated as a percentage of the OD of the treated cultures in comparison with the untreated controls (viability control). Concentrations that maintained more than 90% of cell viability after treatment were considered nontoxic and were used for the antiviral evaluation. Chloroquine (from 6.3 to 100 µM) and Heparin (from 6.3 to 100 µg/mL) were used as positive control of viral inhibition. For the MTT assay, two independent experiments with four replicates each were performed (n=8).

### Evaluation of the antiviral activity

The antiviral activity of ATV against SARS-CoV-2 was evaluated initially through a *pre-post treatment*. Briefly, Vero E6 cells were seeded in 96-well plates with a density of 1.0 x 10^4^ cells/well. Cells were incubated for 24h, at 37°C, 5% CO_2_, and then pretreated with double dilutions of ATV (3.9 – 31.2 µM) for 1h prior to infection. Treatment was then removed, and cells were infected with SARS-CoV-2 stock at a MOI of 0.01 in DMEM with 2% FBS. Cultures were incubated for 1h at 37°C. The inoculum was removed and replaced newly by the same dilutions of the compound. After 48h of treatment, the cell culture supernatants were harvested and stored at −80 °C to be quantified by plaque assay. The supernatant of infected cells without treatment was used as the untreated control.

Additionally, two antiviral strategies were done: *pre-infection treatment* (ATV was added 1h before infection and was removed prior to viral infection) and *post-infection treatment* (ATV was added 48h after infection). Chloroquine (from 12.5 to 100 µM) and Heparin (12.5 −100 µg/mL) were included as positive controls for viral inhibition. Two independent experiments with four replicates per experiment were performed (n=8).

### Viral quantification

The reduction of SARS-CoV-2 viral titer in cell supernatants was quantified by plaque assay. Briefly, tenfold serial dilutions of the supernatants obtained from the antiviral assay were prepared in DMEM with 2% FBS and used to inoculate confluent monolayers of Vero E6 cells into plates of 24 wells (1.0×10^5^ cells/well). After 1h of incubation, viral inoculum was removed, and cells were overlaid with 1.5% carboxymethyl-cellulose in DMEM with 2% FBS. Then, the cultures were incubated for 3 days at 37°C, with 5% CO_2_. After incubation, the monolayers were washed twice with PBS, fixed with 4% Formaldehyde, and plaques were revealed with 1% Crystal violet solution (Sigma-Aldrich). Plaques were counted and used to calculate the number of plaque-forming units per milliliter (PFU/mL). The reduction in the viral titer after treatment compared to the infection control (untreated infected cells) was expressed as inhibition percentage. Two independent experiments with two replicates per each were performed (n=4).

### Molecular Docking

The simulation of molecular docking was used to measure the binding affinity of the chemical functional groups of the ligands (Atorvastatin and Chloroquine) with amino acids in the binding sites of three SARS-CoV-2 proteins.

The selected proteins were considered and obtained from the Protein Data Bank (PDB) [48]. SARS-CoV-2 Spike protein (PDB: 6VYB) [49], RdRp (PDB: 6M71) [50] and 3CL protease (PDB: 6M2N) [51] were selected. The protein structures were subjected to preparation at Discovery Studio [52] and AutoDock Tools (ADT). The ligands were drawn and optimized using Avogadro software [53] and ADT.

PrankWeb [54] was used to check the binding site coordinate and specify the amino acids in the SARS-CoV-2 proteins and pockets. Couplings were carried out using Auto Dock vina version 4.2.6 [55], configuring a box with dimensions x: 24, y: 24, z: 24, this for each simulation carried out with the viral proteins using exhaustiveness of 20. For 3CL protease (PDB:6M2N), coordinates x: −47.585, y: 1.135 z: −5.600; RdRp (PDB:6M71), x:125.507, y:138.079, z: 130.430 and Spike protein (PDB:6VYB), x:210.913 y:211.760, z: 185.437 coordinates were used. Then, we used BIOVIA Discovery Studio Visualizer 16.1 to illustrate 2D docked visualization analysis of the ligands with the amino acids inside the pockets [56].

### Statistical analysis

All data were analyzed with GraphPad Prism (La Jolla, CA, USA) and presented as mean ± SEM. Statistical differences were evaluated via Student’s t-test or Mann–Whitney U test, according to the normality of the data. A value of p ≤ 0.05 was considered significant. The median effective concentration (EC50) values represent the concentration of each medicament that reduces viral titer by 50%. The CC50 values represent the concentration that cause 50% cytotoxicity. The corresponding dose-response curves were fitted by non-linear regression analysis using a sigmoidal model. The calculated selectivity index (SI) represents the ratio of CC50 to EC50.

## Supplementary Figures

**Supplementary Figure 1.**
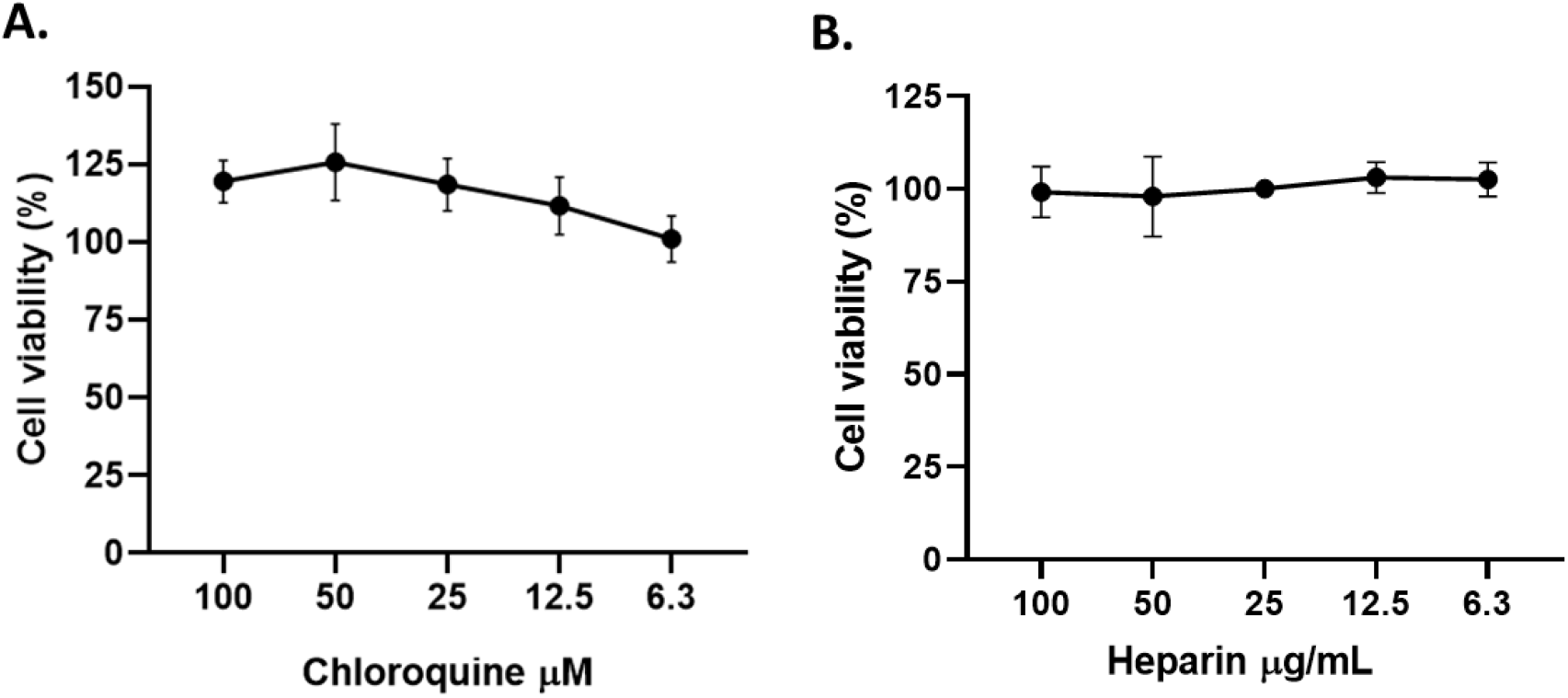
Chloroquine and Heparin did not affect the viability of Vero E6 cells. The figure represents the viability percentage of Vero E6 cells after 48h of treatment with CQ **(A)** and Heparin **(B).** Bars represent mean values ± SEM. Two independent experiments with four replicates each experiment was performed (n=8).

**Supplementary Figure 2.**
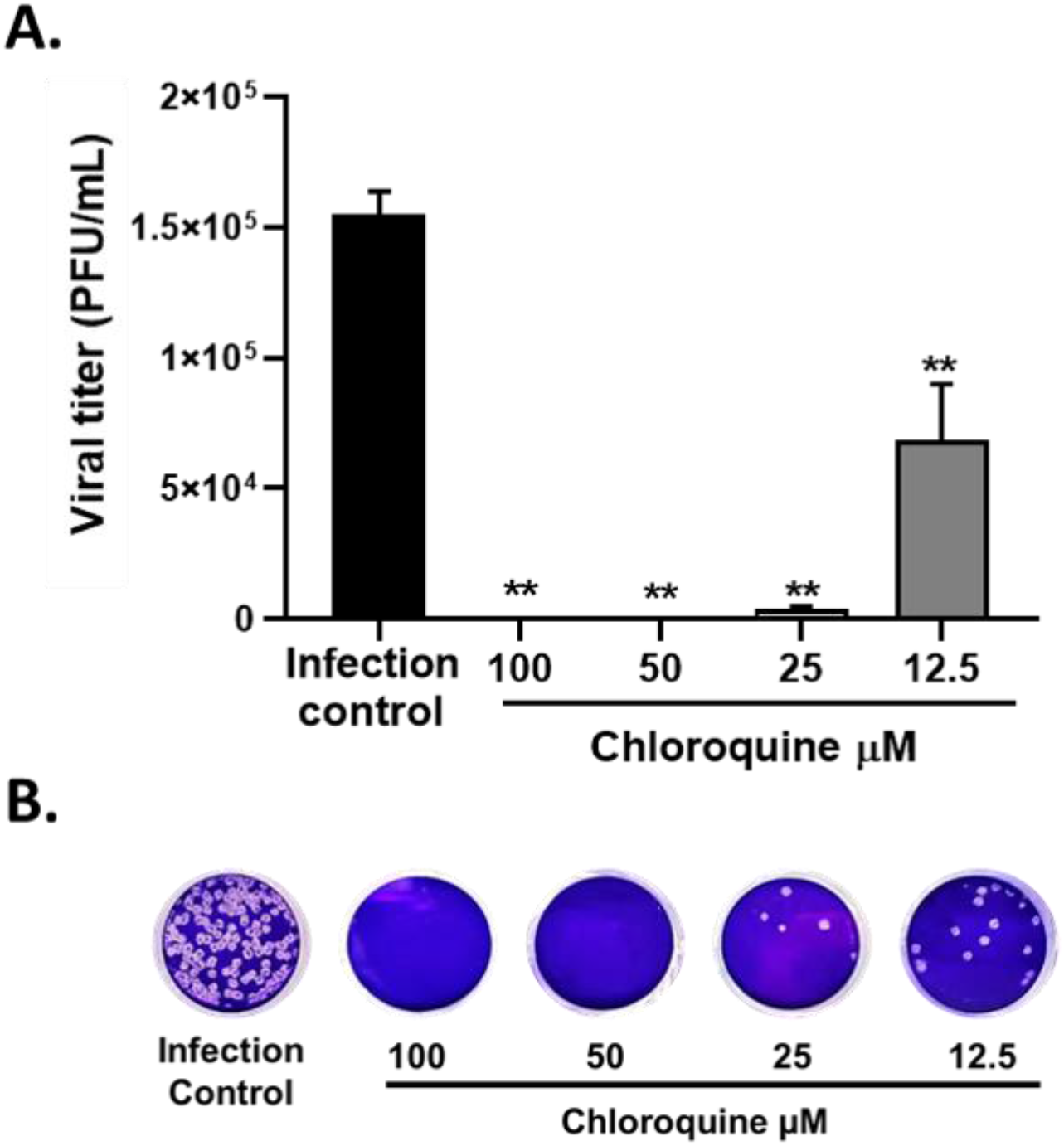
Antiviral activity of Chloroquine against SARS-CoV-2 by pre-post treatment. **A**. The figure represents the viral titer (PFU/mL) of supernatants harvested after the pre-post treatment with CQ, quantified by plaque assay (n=4). Bars represent mean values ± SEM. *p ≤ 0.05, ** p ≤ 0.01. Two independent experiments with two replicates each experiment were performed, n=4. **B.** Representative figure of plaque assays on Vero E6 cells of pre-post treatment with CQ against SARS-CoV-2.

**Supplementary Figure 3.**
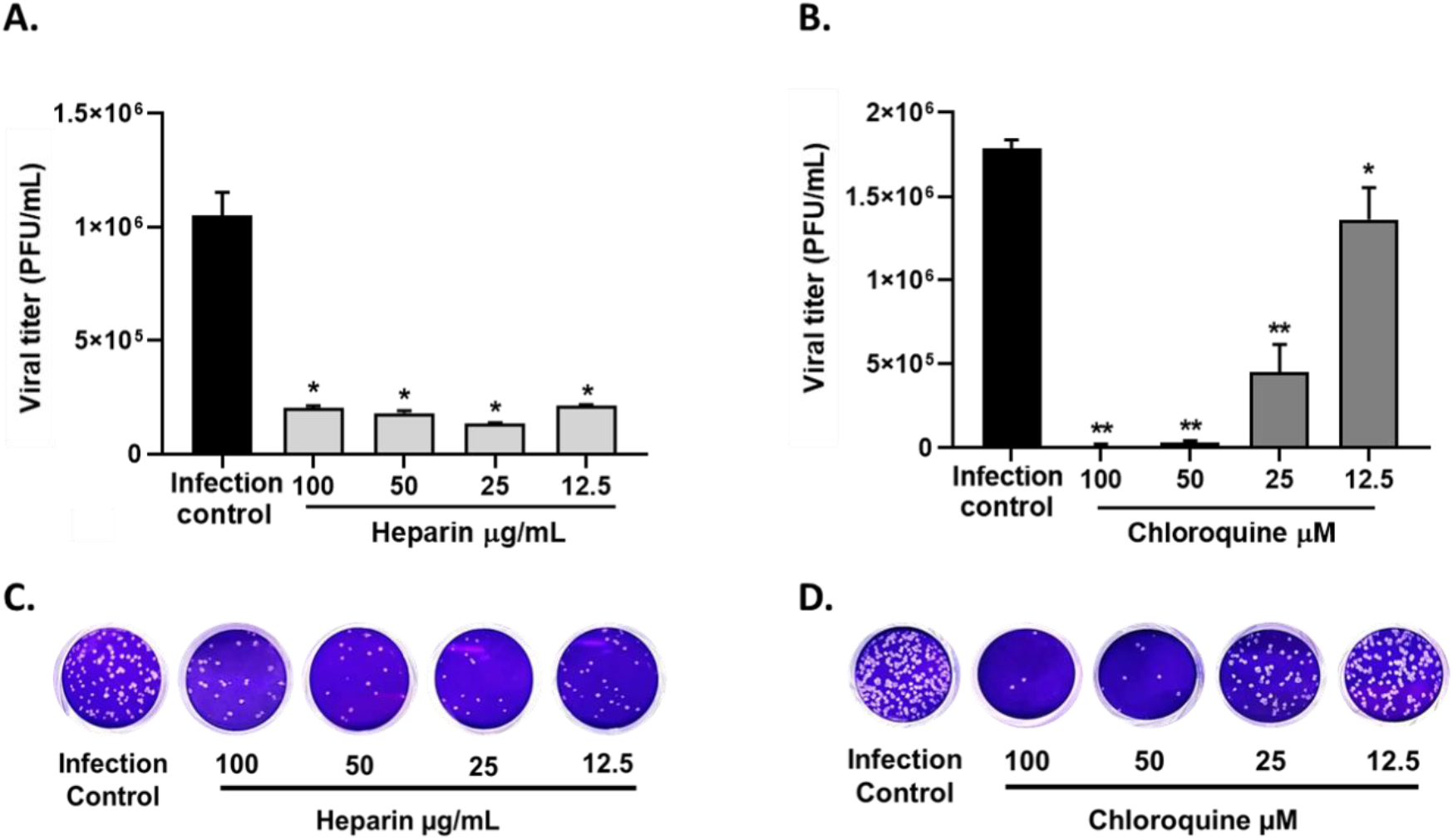
Antiviral activity of inhibition controls against SARS-CoV-2 by pre-infection and post-treatment. The figure represents the viral titer (PFU/mL) of supernatants harvested after the treatment with positive control of viral inhibition, quantified by plaque assay (n=4). Bars represent mean values ± SEM. *p ≤ 0.05, ** p ≤ 0.01. **A**. Pre-infection treatment of Heparin. **B.** Post-infection treatment of CQ. **C.** Representative figure of plaque assays on Vero E6 cells of preinfection treatment with Heparin against SARS-CoV-2. **D.** Representative figure of post-infection treatment with CQ against SARS-CoV-2 on Vero E6.

**Supplementary Figure 4.**
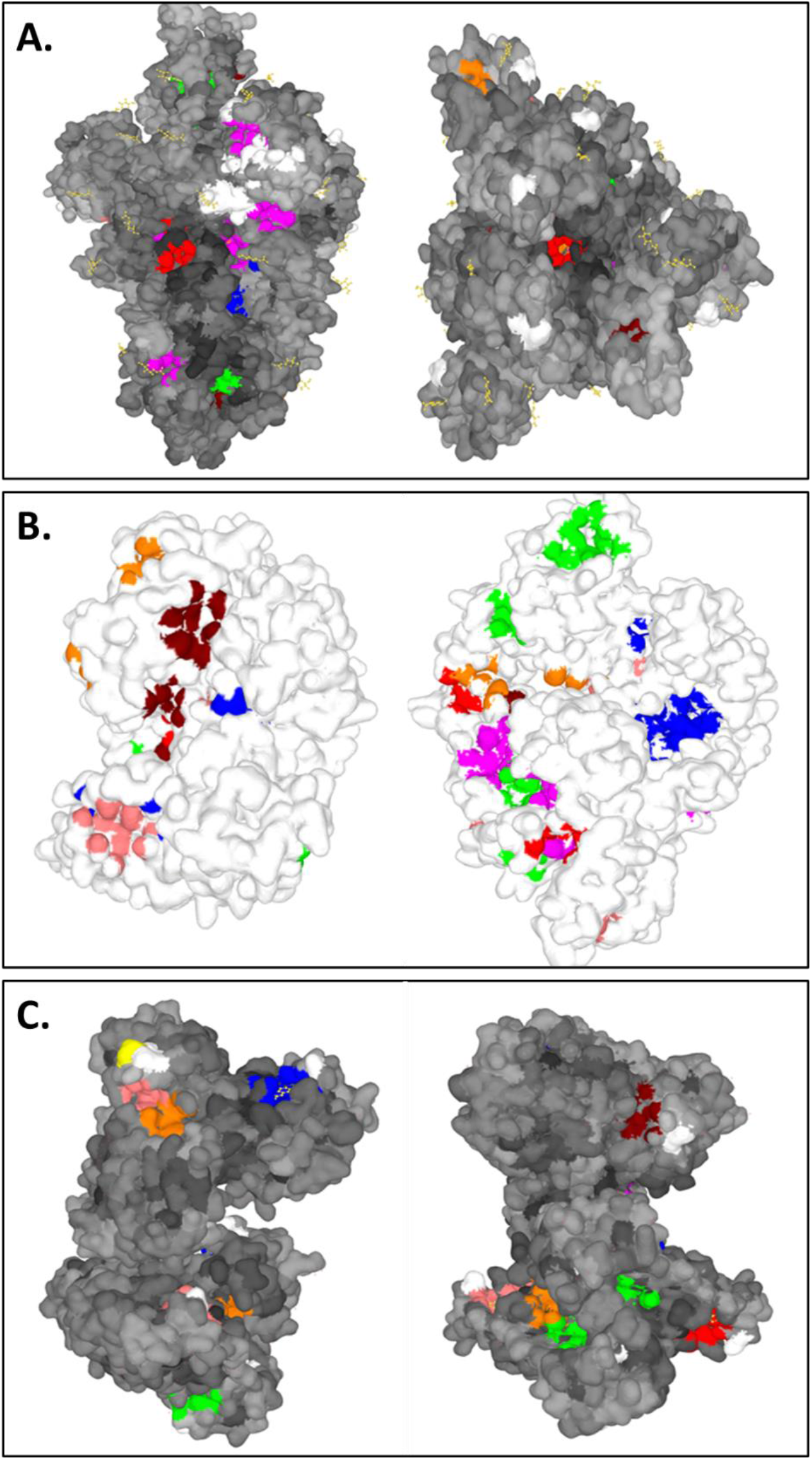
PrankWeb result summary of SARS-CoV-2 proteins, the pockets and the expected amino acids. **A.** Spike protein (PDB:6VYB). **B.** RdRp (PDB: 6M71). **C.** Protease 3CL pro (PDB:6M2N). The first pocket was colored in blue, the second in red, and the third in green.

**Supplementary Figure 5.**
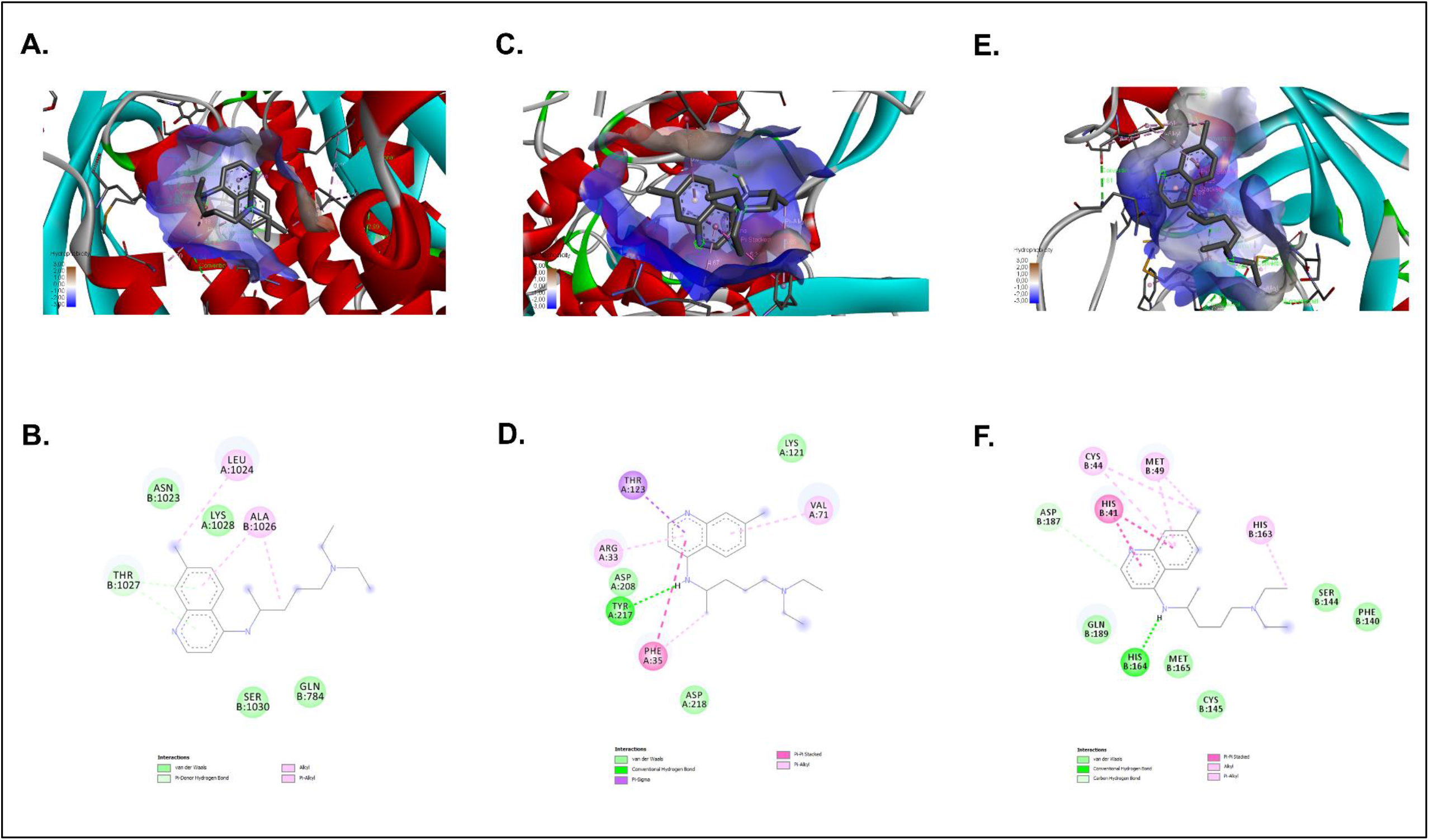
Interaction of CQ with Spike, RdRp and 3CL protease (3CLpro) of SARS-CoV-2. 3D and 2D representations of the main interaction of Chloroquine (positive control of viral inhibition) and three SARS-CoV-2 proteins using BIOVIA Discovery Studio Visualizer 16.1. CQ interaction with: Spike (PDB:6VYB) depicted in 3D **(A)** and 2D **(B)**; RdRp (PDB:6M71) represented in 3D **(C)** and 2D **(D)**; 3CL protease (PDB: 6M2N) of SARS-CoV-2 depicted in 3D **(E)** and 2D **(F).** The interactions formed in the complexes are described in each figure.

## Acknowledgments

We want to thank to Instituto Nacional de Salud, Bogotá-Colombia for donating the Vero E6 cell line.

## DECLARATIONS

### Ethics approval and consent to participate

This study was carried out keeping good records, practicing good data collection and management, transparency of data-sharing and realistic representation of study results.

### Competing interests

None of the authors has any potential financial conflict of interest related to this manuscript.

### Funding

This study was supported by Universidad de Antioquia, CODI and Universidad Cooperativa de Colombia. BPIN 2020000100131-SGR.

### Authors’ Contributions

María T. Rugeles, Wildeman Zapata-Builes, Ariadna L. Guerra-Sandoval, Carlos M. Guerra-Almonacid, Jaime Hincapié-García and Juan C. Hernandez conceived and designed the study. María I. Zapata-Cardona, Lizdany Flórez-Álvarez analyzed the data, interpreted the results and wrote the manuscript. María T. Rugeles, Wildeman Zapata-Builes, Ariadna L. Guerra-Sandoval, Carlos M. Guerra-Almonacid, Jaime Hincapié-García and Juan C. Hernandez assisted with interpretation of the results and manuscript writing and María T. Rugeles guided and reviewed the research. All authors have contributed to editing this paper; they have approved this final submission and declare no conflict of interest.

